# Identification of genes associated with productivity traits and salinity tolerance from activation tagged lines of rice

**DOI:** 10.1101/2021.01.22.427785

**Authors:** Kota Vamsee Raja, Kalva Madhanasekhar, Vudem Dashavantha Reddy, Attipalli Ramachandra Reddy, Khareedu Venkateswara Rao

**Affiliations:** Centre for Plant Molecular Biology, Osmania University, Hyderabad, 500 007, India; Department of Plant Sciences, University of Hyderabad, Hyderabad, 500 046, India

**Keywords:** Activation tagging, Agronomic traits, Crop improvement, Growth and Development, Overexpression of genes, Salinity tolerance, Water use efficiency

## Abstract

World-wide crop productivity is hugely impacted by diverse eco-environmental conditions. In the present investigation, activation tagged (AT) lines of rice endowed with improved agronomic attributes have been analyzed for tolerance to salinity stress besides identification of genes associated with these attributes. Under salinity stress conditions, AT lines exhibited increased seed germination rates, improved plant growth and development at vegetative and reproductive stages as compared to wild-type (WT) plants. Furthermore, AT lines disclosed enhanced plant water content, photosynthetic efficiency, stomatal conductance, water use efficiency and maximum quantum yield when compared to WT plants, leading to improved yields and delayed onset of stress symptoms. Moreover, AT lines revealed effective antioxidant systems causing decreased accumulation of reactive oxygen species and delayed salinity stress symptoms compared to WT plants. Reduced accumulation of malondialdehyde with concomitant increases in proline and soluble sugars of AT lines further endorsing their improved stress tolerance levels. TAIL and qRT-PCR analyses of AT lines revealed Ds element integrations at different loci and respective overexpression of identified candidate genes involved in various aspects of plant development and stress tolerance. Accordingly, the AT lines plausibly serve as a rare genetic resource for fortifying stress tolerance and productivity traits of elite rice cultivars.

**Highlight:** Activation tagged lines of rice endowed with improved agronomic attributes have been analyzed for tolerance to salinity stress besides identification and expression analysis of genes associated with these attributes.

## Introduction

Globally, diverse cereal crops serve as a major source of food and essential nutrients for humans and also as feed for the livestock and domestic animals. With ever increasing global populations, the demand for food has been rising rapidly. Whereas, on the other hand, the crop productivity is impacted severely and hampered by various eco-environmental factors, viz., global warming, abiotic, biotic stresses, etc., posing serious threats to the world food security (Maiti and Satya, 2014). Rice (*Oryza sativa*, L) serves as the foremost staple food for more than half of the world’s population. Numerous abiotic stresses have been found to cause huge (>70%) damage to the rice crop during its growth and development (Akram *et al*., 2019). Moreover, these abiotic stress effects proved to be even more severe in conjunction with other environmental constrains such as air pollution, nutrition limitations and biotic factors, resulting in further reductions in crop yields triggering social unrest and famines (Xu *et al*., 2015; Sekhar *et al*., 2020). For ensuring food and nutritional security, it is essential to tailor make resilient crops for addressing the problems of diverse abiotic stress effects through genetic engineering and molecular breeding approaches (Zafar *et al*., 2020; Nutan *et al*., 2020).

Among the various abiotic stresses, drought and salinity have been found to impose drastic adverse effects on yields and productivity of diverse crop plants (Munns and Tester, 2008; Pasala *et al*., 2016). Globally, salinity was found to affect about 33% of the irrigated agricultural lands amounting to 20% of the total cultivated lands (Jamil *et al*., 2011). In plants, salinity induces osmotic stress and reduce water uptake resulting in decreased relative water content and water potential thereby causing retarded growth, stomatal closure, declined photosynthesis and transpiration. Reduced crop yields under salinity have been mainly due to diminished photosynthetic efficiency caused by stomatal or non-stomatal limitations under varied stress intensity, its duration and plant species (Gupta and Huang, 2014; Shahzad *et al*., 2019; Pan *et al*., 2020).

Crop cultivars grown under salinity conditions were imposed with both osmotic and ion-specific stresses causing overproduction of reactive oxygen species (ROS). For scavenging the excess generated ROS, plants could upregulate the production of low molecular mass antioxidants, viz., ascorbic acid, reduced glutathione, besides antioxidant enzymes such as superoxide dismutase, catalases, ascorbate peroxidases and glutathione peroxidases (Bartwal *et al*., 2013). Oxidative stress tolerance was induced in plants by regulating antioxidant enzyme activities and by coordinating the action of numerous multiple stress responsive genes which interact with diverse components of stress signal transduction pathways (Hasanuzzaman *et al*., 2012).

Tolerance to salinity stress was found to be a multigenic attribute and hence identification of new genes conferring salt tolerance has become essential for the improvement of rice productivity (Chinnusamy *et al*., 2005). Furthermore, rice crop yield was reported to be associated with the rate of photosynthesis, amount of light incident on leaves and the atmospheric carbon dioxide captivated by the leaves during photosynthesis (Ambavaram *et al*., 2014). As such, it is essential for the plant to coordinate and synergize the mechanisms of leaf gas exchange, photosynthesis and other physiological processes to promote improved crop yields under different stress conditions. Numerous agronomic and physiological characters contributing to improved grain yields include plant height, total biomass, spike length, number of grains per panicle, water-soluble carbohydrates, chlorophyll content besides genetic makeup (C3/C4) and efficiency of photosynthesis (Gao *et al*., 2017). Moreover, genetic advance in crop productivity potential can be attained by introgressing specific genes governing improved agronomic attributes as well as physiological performance under various biotic and abiotic stresses (Lopes *et al*., 2012). Further, for developing superior quality crops, it is imperative to identify and analyze the functionality of diverse genes responsible for improved performance and increased productivity under different stress conditions. Hsing *et al*. (2007) reported that generation of a high frequency of activation tagged (AT) mutants in crops serves as a rich genetic resource for analyzing various candidate genes of agronomic importance.

Earlier, we generated a large population of AT mutants in the rice cultivar BPT 5204, adopting activation tagging approach (Kota *et al*., 2020). In the present investigation, from among diverse mutant lines generated, select mutant rice lines exhibiting marked increments in flag leaf length, plant height, tiller number per plant, panicle length, number of grains per plant and grain length have been chosen for screening their potential ability to withstand the effects of salinity stress. Furthermore, we have analyzed the various changes in the processes underlying physiological and antioxidant metabolisms as compared to the wild type (WT) plants under salinity stress conditions at different stages of plant growth and development. Additionally, we have also characterized the flanking sequences of integrated *Ds* element in the genomes of AT lines and determined expression levels of activated genes.

## Materials and methods

### Genetic transformation and generation of AT mutants

Activation tagging vector pSQ5 was obtained from Prof. Venkatesan Sundaresan, University of California, Davis, USA, and was employed for the development of AT mutants in an indica rice cv. of Samba Mahsuri (BPT 5204). A total number of 10 primary transformants, as confirmed by PCR and Southern blot analyses, have been used for generation of more than 6000 stable *Ds-RFP* AT lines (Kota *et al*., 2020). Based on phenotypic variations exhibited by AT mutants, three identified lines of A10-*Ds-RFP*6, F4-*Ds-RFP*31 and T8-*Ds-RFP*3 have been selected for further analyses and characterization.

### PCR analysis of AT lines

Analyses of A10-*Ds-RFP*6, F4-*Ds-RFP*31 and T8-*Ds-RFP*3 AT lines were carried out employing the primers of *RFP* forward: 5’-AGGACGGCTGCTTCATCTAC-3’ & *nos* reverse: 5’-TTGCGCGCTATATTTTGTTTT-3’, and enhancer forward: 5’-CAAAGGGTAATAT CGGGAAACC-3’ & enhancer reverse: 5’-TCACATCAATCCACTTGCTT-3’ to amplify *RFP* and 4X enhancer regions, respectively. DNA isolated from wild type (WT) plant was used as the negative control and intermediate vector was used as the positive control.

### Salinity stress treatments at different developmental stages of AT lines Seed germination assays

Seeds of AT lines A10-*Ds-RFP*6, F4-*Ds-RFP*31 and T8-*Ds-RFP*3 along with WT were surface sterilized with 0.1% (w/v) HgCl_2_ for 8 min followed by washing thrice with autoclaved distilled water. The sterilized seeds were then transferred onto the ½ MS medium (control) and media containing 250 mM NaCl for salinity stress treatments. Data on germination frequencies were recorded after 14 days of stress treatments. Seed germination rate (%) was computed using the formula:

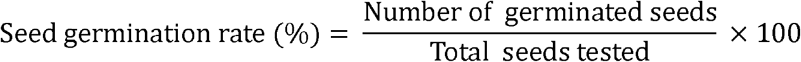

### Stress treatments at the seedling stage

To test the salinity tolerance ability of AT lines during the initial growth stage, seedlings were grown in Hoagland solution with a 16-h light/8-h dark cycle at 30°C in the glass house along with WT plants. The seedlings of 15 days-old were treated with 250 mM NaCl for salinity stress for 10 days. After 7 days of recovery under normal conditions, the seedlings were photographed and data on survival rate, biomass, shoot and root lengths were recorded.

### Stress treatments at the vegetative stage of plants

For stress treatment at the vegetative stage, AT lines and WT plants were grown in pots at 30°C in the glass house. Plants at the age of 60-65 days old were then subjected to 250 mM NaCl stress for 15 days. After stress treatments, the NaCl solution was replaced with fresh water and plants were allowed to grow to maturity under normal conditions. Data on photosynthetic leaf gas exchange parameters, oxidative stress parameters, osmolytes, antioxidants, antioxidant enzymes and leaf relative water content (RWC) were analyzed.

### Stress treatments at the reproductive stage of plants

For stress experiments at the reproductive stage, AT lines and WT plants were grown in pots in the glass house. Salinity (250 mM NaCl) stress was applied to AT lines (90-100 days old) along with WT plants for 15 days. After stress treatments, the plants were allowed to grow to maturity under normal conditions. Data on panicle length and number of grains per plant were recorded and analyzed.

### Leaf relative water content, photosynthetic leaf gas exchange characteristics and maximum quantum yield

To check the leaf RWC, fresh leaf discs (1.5 cm^2^) sampled from A10-*Ds-RFP*6, F4-*Ds-RFP*31 and T8-*Ds-RFP*3 lines, subjected to well-water and salinity stress along with WT plants, were weighed immediately (fresh weight). A set of leaf discs were immersed in water and incubated at 4°C for 24 h (desiccation weight) and a second set of leaf discs were dried at 70°C for 24 h simultaneously and dry weights were recorded. The relative water content was estimated using the formula of Barrs and Weatherley (1962).

*In-situ* photosynthetic measurements were carried out between 9:30 AM to 11:30 AM using a portable infrared CO_2_/H_2_O gas analyzer (IRGA) LI-6400XT (LI-COR Inc., NE, USA). The following conditions were maintained during the measurements: a saturating photosynthetically active radiation (PAR) of 1600 μmol m^-2^ s^-1^, air temperature of 25–26 °C, and relative humidity of 55–60%. Photosynthetic leaf gas exchange measurements of A10-*Ds-RFP*6, F4-*Ds-RFP*31 and T8-*Ds-RFP*3 AT mutants along with WT plants were recorded at ambient [CO_2_] (420 μmol mol^-1^) conditions using young, well expanded, light exposed and randomly chosen leaves under well-water and salinity stress conditions. The leaf from selected plant was enclosed in the leaf chamber for two minutes. During this time the leaf gets adapted to the new microclimate; later data on the light saturated net photosynthetic rates (A_Sat_; μmol m^-2^ s^-1^), stomatal conductance (*g_s_*; mol m^-2^ s^-1^), transpiration rates (*E*; mmol m^-2^ s^-1^) and intercellular [CO_2_] (C_i_; μmol) were recorded. Furthermore, data pertaining to light saturated net photosynthetic rates and transpiration rates were used to calculate the instant water use efficiency (WUE_i_=A_Sat_/E; mmol CO_2_ mol^-1^ H_2_O). In addition, light (1600 μmol m^-2^ s^-1^) and CO_2_ (1200 μmol m^-2^ s^-1^) saturated photosynthetic rates (A_Max_) were measured in all the representative samples.

Further, chlorophyll a (Chl a) fluorescence measurements were taken on the same leaves used for the photosynthetic leaf gas exchange measurements by using MINI-version of the Imaging PAM (HeinzWalz GmbH, Effeltrich, Germany) on the adaxial side of the leaves with saturated light flashes. Maximal photochemical efficiency of photosystem-II, (F_M_–F_0_)/F_M_ = F_V_/F_M_, was calculated from the leaves which were pre-dark adapted for 30 min in both AT lines and WT plants under well-water and NaCl stress conditions.

### Estimation of total chlorophyll content and reducing sugars

The leaf discs of 0.53 cm^2^ collected from well-water and NaCl stressed AT lines along with WT plants were ground into fine powder in liquid nitrogen and mixed with 80% acetone. After thorough mixing, the supernatant was collected by centrifuging the tubes at 12,000 rpm for 15 min. Total chlorophyll content was estimated according to Lichtenthaler and Wellburn (1983).

Furthermore, leaf samples of 100 mg were collected from well-water and salinity treated AT lines as well as WT plants to quantify the reducing sugars. The leaf tissue was immediately frozen and ground to fine powder in liquid nitrogen. From leaf powder samples, reducing sugars were extracted using 80% ethanol at 95°C twice. The supernatant was concentrated by subjecting to drying at 80°C for 2 h. The sample extracts were prepared by dissolving the residue in 10 ml of distilled water. Finally, reducing sugar content values were estimated as per the method described by Miller (1959) and expressed in mg g^-1^ FW.

### Quantification of oxidative stress parameters, osmolytes and antioxidants

In order to assess the oxidative stress effects, lipid peroxidation was determined by quantifying malondialdehyde (MDA) content. About 100 mg of leaves were collected from well-water and salinity treated AT lines along with WT plants, which were homogenized in 10 ml of 10% trichloroacetic acid and centrifuged at 5000 g for 10 min. The MDA content in the supernatant was quantified by measuring it using the thiobarbituric acid reaction as described by Fu and Huang (2001) and expressed in μmol g^-1^ FW.

Similarly, stress induced osmolyte, proline content was estimated in AT lines as well as WT plants under well-water and salinity stress conditions. Fresh leaf samples (0.5 g) of AT mutants, subjected to well-water and salinity stress along with WT plants, were homogenized in 5 ml of 3% sulfosalicylic acid and incubated in a boiling water bath for 10 min. After incubation, the tubes were centrifuged at 10,000 rpm for 10 min at 4°C and the supernatant was used for estimation of proline content as per Bates *et al*. (1973), from the standard curve and expressed in μg g^-1^ FW.

Non-enzymatic antioxidants including ascorbic acid and total phenolic contents were also measured. Fresh 0.5 g leaf samples collected from AT lines and WT plants, subjected to well-water and salinity stress, were homogenized in 5 ml of 10% trichloroacetic acid and were centrifuged at 10,000 g for 20 min at room temperature. Ascorbic acid (ASA) present in the leaf extracts was estimated adopting the protocol of Omaye *et al*. (1979). The ASA content was determined from the standard curve and expressed in mg g^-1^ FW. Total phenolic contents were estimated in AT mutants as well as WT plants after subjecting them to well-water and salinity stress, according to the method of Stankovic (2011) and were expressed in gallic acid equivalents as mg of GA g^-1^ FW.

### Estimation of antioxidant enzyme activities

Antioxidant enzyme activities of ascorbate peroxidase (APX), glutathione peroxidase (GPX), monodehydroascorbate reductase (MDHAR), glutathione reductase (GR), superoxide dismutase (SOD) and catalase (CAT), were estimated in AT lines and WT plants following the protocols of Murshed *et al*. (2008) with minor modifications. The leaf samples (0.5 g) collected were finely powdered in liquid nitrogen and mixed with 2.5 ml of 50 mM potassium phosphate buffer containing 0.04 M KCl, 1 mM reduced ascorbate (ASC), 1 mM PMSF and 8% PVPP (pH 6.0). The homogenates were centrifuged at 14,000 g for 10 min at 4°C and the supernatants were used for estimating the enzyme activities. The APX activity was measured in the leaf tissues of AT lines and WT plants according to the method of Nakano and Asada (1981) by monitoring the rate of ascorbate oxidation at 290 nm and APX activity was expressed in terms of μmol min^-1^ g^-1^ FW. Further, GPX activity of AT mutants along with WT plants was estimated according to the protocol of Paglia and Valentine (1967). The final GPX enzyme activity was calculated using extinction coefficient 6.22 mM^-1^ cm^-1^ and expressed in terms of mmol NADPH mg protein^-1^ min^-1^. The MDHAR activity was quantified as per the protocol described by Miyake and Asada (1992) employing the leaf samples of AT lines and WT plants. Specific activity of the enzyme was estimated using the extinction coefficient of 6.22 mM^-1^ cm^-1^; and MDHAR activity was expressed in terms of nmol min^-1^ g^-1^ FW. To estimate the GR activity in AT lines and WT plants, 5 μl of 20 mM GSSG was added to 1 ml reaction mixture containing 50 mM phosphate buffer, 10 mM EDTA and 25 μl of sample supernatant. GR activity was assayed spectrophotometrically based on the NADPH oxidation at 340 nm in the presence of 0.2 mM NADPH. The specific GR activity was calculated using 6.22 mM^-1^ cm^-1^ extinction coefficient and expressed as nmol min^-1^ g^-1^ FW. The SOD activity of AT lines and WT plants was estimated by monitoring the inhibition of blue formazane production by means of photochemical reduction of nitro blue tetrazolium (NBT) according to Beauchamp and Fridovich (1971). One unit of SOD activity was defined as the amount of enzyme required to inhibit 50% of the NBT photo reduction and is expressed as U mg protein^-1^ min^-1^. Catalase activity of AT lines and WT plants was determined as described by Aebi (1984). The enzyme activity in leaf tissues was calculated using the molar extinction coefficient of H_2_O_2_ (36 mM^-1^ cm^-1^) and expressed as moles of H_2_O_2_ consumed min^-1^ mg^-1^ protein.

### Thermal Asymmetric Interlaced PCR (TAIL-PCR) analysis

TAIL-PCR analysis was performed as per the conditions described by Liu and Chen (2007) with appropriate modifications. The primary TAIL-PCR amplification was performed using reaction (20 μl) mixture containing 2 μl of 10X PCR buffer, 200 μM of each dATP, dCTP, dGTP and dTTP, 0.3 μM of 3’ *Ds*-1 specific primer (Kolesnik *et al*., 2004), 5 μM AD2 degenerate primers (Liu and Chen, 2007), 0.5 units of Takara Taq polymerase (Takara Bio Inc., Japan) and 50 ng genomic DNA of selected AT mutants. Secondary TAIL-PCR reaction was carried out in 20 μl volume containing 1X Taq buffer, 200 μM dNTPs, 0.2 μM 3’ *Ds*-2 specific primer, 4 μM of the AD2 primer, 0.8 U of Taq polymerase and 50-fold diluted primary PCR product as a template. Tertiary TAIL-PCR was carried out with 50-fold dilution of the secondary PCR product in a reaction mixture containing 1X Taq buffer, 200 μM dNTPs, 0.2 μM 3’ *Ds*-3 specific primer, 3 μM AD2 primer and 0.5 U of Taq polymerase. Thermal cycling conditions adopted for TAIL-PCR are represented in Supplementary Table S1. Tertiary PCR products corresponding to 2° PCR products were excised from the gel, purified and sequenced using the ABI PRISM 3730 DNA Analyzer System (ABI, USA). Flanking sequences were deployed to BLAST against rice genome in the Rice Annotation Project Database (RAP-DB) to check sequence similarity with the rice genome (http://rice.plantbiology.msu.edu/). Results showing high similarity with an E-value <0.01 were taken in to consideration.

### Expression analyses of specific selected genes from AT lines by qRT-PCR

qRT-PCR analyses were performed using the RNA isolated from AT lines and WT plants. First strand cDNA was synthesized from RNA samples and the resultant cDNAs were utilized as templates for qRT-PCR analyses. qRT-PCR was performed using SYBR green master mix with Applied Biosystems 7500 real time PCR system (USA) at 94°C (1 min), 60°C (1 min) and 72°C (1 min) for 30 cycles. The products obtained were analyzed through a melting curve analysis to check the specificity of PCR amplification. Each reaction was performed thrice and relative expression ratios were computed using 2^-ΔΔct^ method employing actin gene as a reference (Livak and Schmittgen, 2001). The primers used for qRT-PCR are listed in Supplementary Table S2.

### Statistical analyses

Data on leaf gas exchange parameters, osmolytes, molecular antioxidants, chlorophyll content, oxidative stress parameters, biomass and yield of AT lines and WT plants subjected to well-water and salinity stress were represented as mean ± SE. In each stress experiment, 10 random plants were used and experiments were repeated thrice. Estimation of mean values, t-test and standard error were carried out using preloaded software in Excel.

## Results

### Development of AT lines of rice

Earlier, we reported generation of a large number of diverse AT mutant lines in rice through *Agrobacterium*-mediated genetic transformation method, employing pSQ5 and pSQ5*bar* vectors containing maize *Ac/Ds* and CaMV35S 4X enhancer elements (Kota *et al*, 2020). Among various AT lines, certain of them exhibited unique distinctive morphological variations, viz., earlier flowering, increases in flag leaf length & breadth, number of tillers, enhanced panicle length, plant height, panicle size, grain colour, increased grain size and grain number/plant (Kota *et al*., 2020). In this report, three AT lines, viz., A10-*Ds-RFP*6 showing increases in tiller number, panicle size and grain number/plant; F4-*Ds-RFP*31 with enhanced flag leaf length & grain length; and T8-*Ds-RFP*3 exhibiting increased root length, tiller number & grain number/plant, obtained with pSQ5 vector, have been chosen mainly to determine the integration site of *Ds* element in their genomes and expression analyses of selected genes present up to 10 kb region on either side of the *Ds* position. Furthermore, an attempt has been made to evaluate their salinity tolerance levels at different developmental stages.

### PCR analyses of AT plants

PCR analyses of three AT lines disclosed a specific 546 bp amplification band corresponding to the *RFP* expression unit with *RFP* forward and *nos* reverse primers (Supplementary Fig. S1) besides four distinct bands of 250 bp, 550 bp, 850 bp and 1150 bp corresponding to the 4X enhancer region (Supplementary Fig. S2) when *En* forward and *En* reverse primers were employed; while WT plants failed to show any similar amplifications.

### Effect of salinity stress at different developmental stages of AT lines

AT rice lines disclosed varied responses against salinity stress at different stages ranging from seed germination to reproductive stage as compared to WT plants. To evaluate the salinity stress tolerance ability of AT mutant lines at seed germination stage, seeds of mutant lines along with that of WT were germinated on MS media supplemented with 250 mM NaCl. After two weeks of salinity stress, AT lines A10-*Ds-RFP*6, F4-*Ds-RFP*31 and T8-*Ds-RFP*3 revealed higher seed germination rates of 78.59 ± 1.59%, 82.61 ± 2.58% and 79.42 ± 2.61% than that of WT (09.94 ± 2.46%) (Fig. 1A). Two week-old seedlings of AT lines subjected to 250 mM salinity stress for ten days exhibited higher survival rates and biomass as well as enhanced shoot and root lengths in comparison to the WT seedlings (Fig. 1B, Fig. 1C, Fig. 1D, Fig. 1E, Fig. 1F). Similarly, the AT plants subjected to 250 mM salinity stress for 15 days at the vegetative stage (55 to 60 days old) reached the reproductive stage and set seeds while WT plants could not reach the reproductive stage (Fig. 2A). Likewise, AT plants, subjected to 250 mM salinity stress for fifteen days at the reproductive stage (90 to 100 days old), developed longer panicles (15.12 ± 0.68 to 16.11 ± 0.61 cm) than that of WT (09.69 ± 0.29 cm) plants (Fig. 2B, Fig. 2C, Fig. 2D). Moreover, AT lines could produce significantly higher number of grains per plant (~ 760 to ~780 seeds) compared with WT plants (~105 seeds) grown under similar stress conditions (Fig. 2B, Fig. 2C, Fig. 2E).

**Fig. 1.**
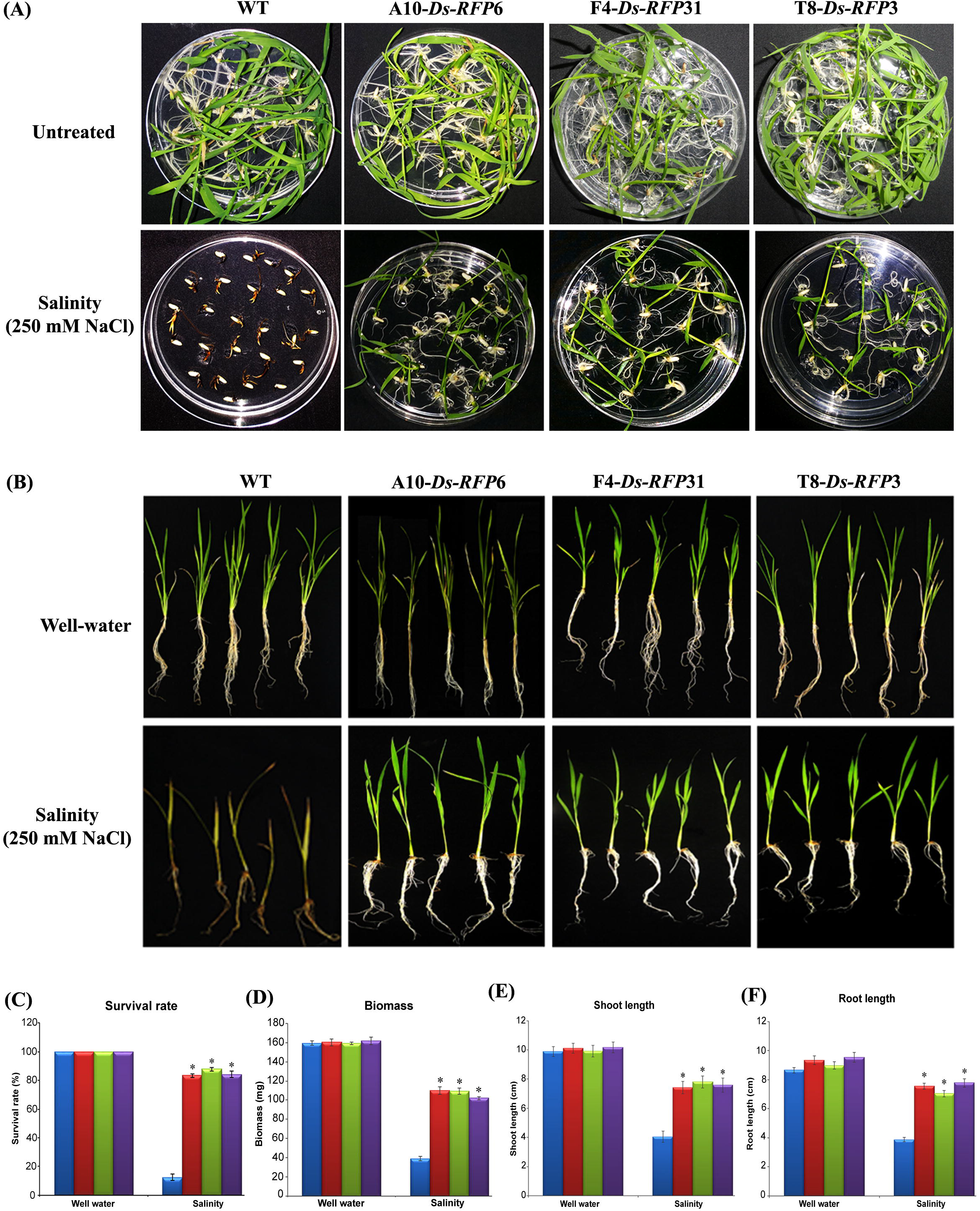
Evaluation of AT rice lines against salinity stress at seed germination and seedling stages. **(A)** Seed germination profiles of WT and AT plants under normal and 250 mM NaCl stress. In each treatment, 20 seeds of WT and three AT lines were used. **(B)** Two-week-old seedlings of WT and AT plants were grown in the presence of 250 mM NaCl for 10 d and were later allowed to recover under normal conditions. Data on survival rate **(C)**, total biomass (**D**), shoot length **(E)** and root length **(F)** were recorded after 7 d of recovery. In each treatment, 10 seedlings of each WT and three AT lines were used. Bar represents mean and error bar represents SE from three independent experiments. * indicates significant differences in comparison with WT at P <0.05. A10-*Ds-RFP6*, F4-*Ds-RFP*31 and T8-*Ds-RFP3* are three different AT lines, WT: Wild type.

**Fig. 2.**
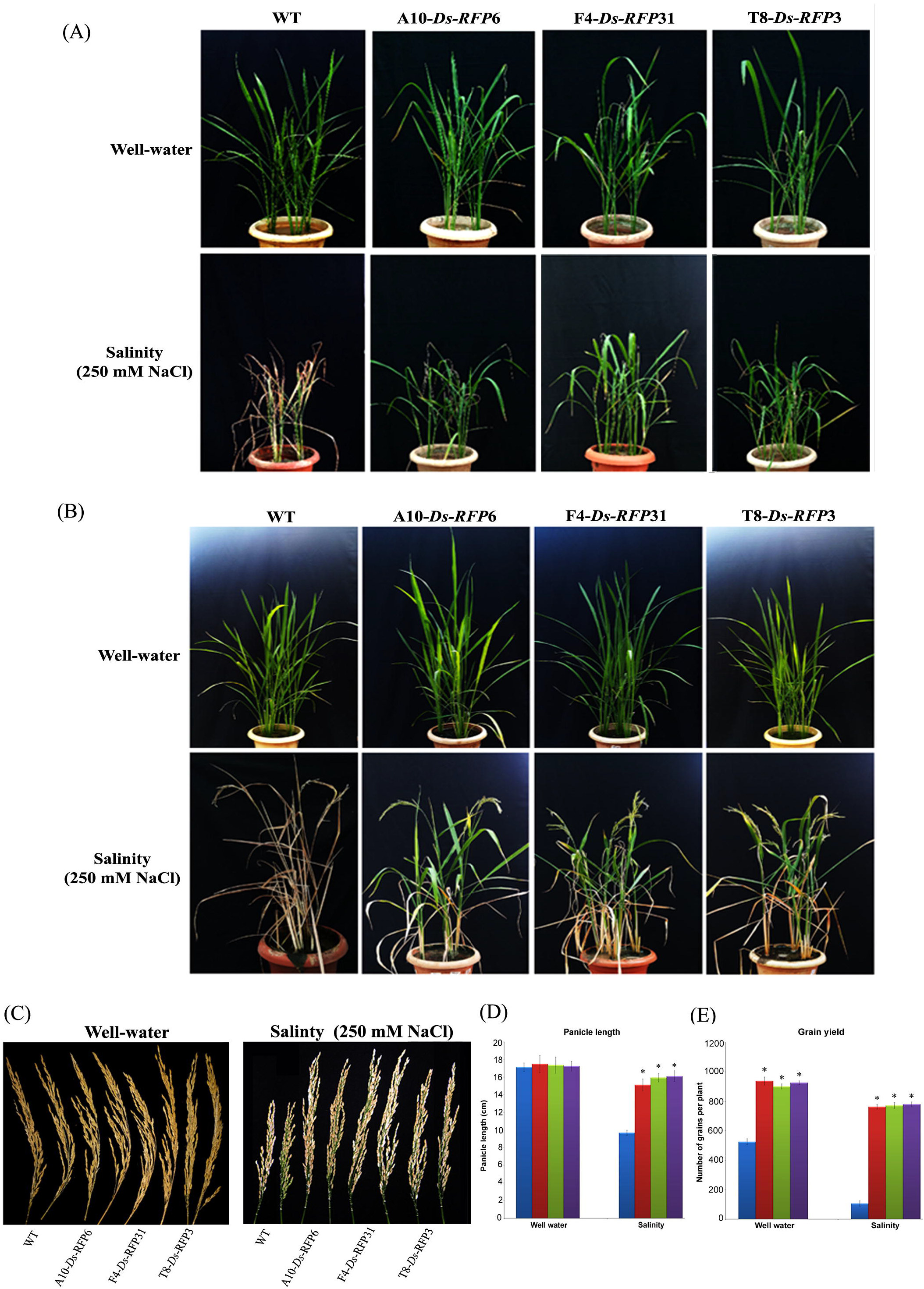
Evaluation of AT mutant lines against salinity stress at vegetative and reproductive stages. AT lines and WT plants were subjected to 250 mM NaCl stress for 15 days at **(A)** vegetative stage (60-65 days old) and **(B)** reproductive stage (90-100 days old). After stress treatment, plants were allowed to grow under normal conditions till maturity. **(C)** Variations of panicle lengths and grain yield of AT lines and WT plants under well-water and salinity stress conditions. Representation of variations in panicle length **(D)** and grain yield per plant **(E)** of AT and WT plants under well-water and salinity stress conditions. In each treatment, 10 plants of each WT and three AT lines were used. Bar represents mean and error bar represents SE from three independent experiments. * indicates significant differences in comparison with WT at P <0.05. A10-*Ds-RFP6*, F4-*Ds-RFP31* and T8-*Ds-RFP3* are three different AT lines, WT: Wild type.

### Leaf relative water content, photosynthetic leaf gas exchange characteristics and maximum quantum yield in AT lines of rice

In addition to morphological improvements, significant variations were also observed in leaf RWC, photosynthetic leaf gas exchange characteristics and maximum quantum yield (F_V_/F_M_) under well-water as well as salinity conditions between the mutants and the WT plants (Fig. 3). However, under well-water conditions, no significant differences were observed in the leaf RWC of mutants and the WT plants. However, AT lines disclosed higher RWC (86 to 87%) under salinity conditions when compared to that of WT (53%) plants (Fig. 3A). The mutants recorded A_Sat_ values ranging from 16 to 21 μmolm^-2^ s^-1^ as compared to 12 μmol m^-2^ s^-1^ of the WT under well-water conditions. Although, imposition of salinity reduced A_Sat_ in both WT (10 μmol m^-2^ s^-1^ to 3 μmol m^-2^ s^-1^) and mutants (15 to 20 μmol m^-2^ s^-1^ to 1014 μmol m^-2^ s^-1^), yet the reduction was more in the WT plants when compared to that of AT lines. These results indicate that AT lines showed higher A_Sat_ values by 25% to 50 % under well-water and by 50% to 90% under salinity stress conditions when compared to the WT plants (Fig. 3B). With respect to A_Sat_, the AT lines exhibited increases in stomatal conductance (g_s_) under both well-water as well as salinity conditions when compared to the WT plants. The observed g_s_ values in mutants ranged from 0.239 to 0.317 mol m^-2^ s^-1^, while it was 0.162 mol m^-2^ s^-1^ in the WT plants. Further, imposition of salinity stress reduced g_s_ values (0.165 to 0.198 mol m^-2^ s^-1^) in AT lines and WT plants (0.096 mol m^-2^ s^-1^) (Fig. 3C). Likewise, alterations in the transpiration (E) rates showed similar trend as that of g_s_ under well-water and salinity conditions in AT and WT plants. The observed E in WT plants under well-water conditions was 3.8 mmol m^-2^ s^-1^, while in the AT lines it varied from 4.9 to 5.8 mmol m^-2^ s^-1^. Imposition of salinity stress reduced E values in WT plants as well as mutants, yet the reduction in E was higher in the WT plants when compared to that of mutants (Fig. 3D). Whereas, for photosynthetic leaf gas exchange parameters, non-significant differences were noticed in intercellular CO_2_ (C_i_) concentrations between WT plants and AT mutants under both well-water (239 to 247 μmol) and salinity stress (189 to 208 μmol) conditions (Fig. 3E). Moreover, when AT mutants were tested for their instant water use efficiency (WUE_i_, A_Sat_/E) under various representative conditions, results revealed significant differences among them. Under well-water conditions, the WUE; in WT plants was 2.76 mmol CO_2_ mol^-1^ H_2_O, while in the AT mutants it varied from 3.01 to 3.51 mmol CO_2_ mol^-1^ H_2_O. Imposition of salinity stress on WT plants caused marked reduction in WUE_i_ from 2.76 to 1.98 mmol CO_2_ mol^-1^ H_2_O; whereas, AT lines revealed significantly higher WUE_;_ values ranging from 3.28 to 5.11 mmol CO_2_ mol^-1^ H_2_O (150% to 250%) in contrast to WT plants (Fig. 3F). Moreover, both light and CO_2_ saturated photosynthetic rates (A_MAX_) were significantly higher in the mutants both under well-water and salinity stress conditions when compared to WT plants. Under well-water conditions, the observed A_MAX_ in WT plants was 29.05 μmol m^-2^ s^-1^, while in the mutants it ranged from 30.87 to 32.36 μmol m^-2^ s^-1^. Even though, imposition of salinity stress could reduce A_MAX_ in WT as well as among mutant lines, yet the level of reduction was higher in WT plants as compared to the mutants. In WT plants, salinity stress reduced A_MAX_ from 29.05 to 12.05 μmol m^-2^ s^-^, while in mutant lines the value was reduced from 30.87-32.02 μmol m^-2^ s^-1^ to 20.84-21.65 μmol m^-2^ s^-1^ (Fig. 3G). Under well-water conditions, AT lines, compared to WT plants, did not show significant differences between them in their maximum quantum yields (F_V_/F_M_). However, imposition of salinity stress reduced F_V_/F_M_ in both AT lines and WT plants while the reduction was higher in the WT plants. The observed F_V_/F_M_ values in WT plants under well-water and salinity stress conditions were 0.74 and 0.55, respectively; while F_V_/F_M_ of AT lines under identical conditions varied from 0.77 to 0.79 and 0.68 to 0.72, respectively (Fig. 3H).

**Fig. 3.**
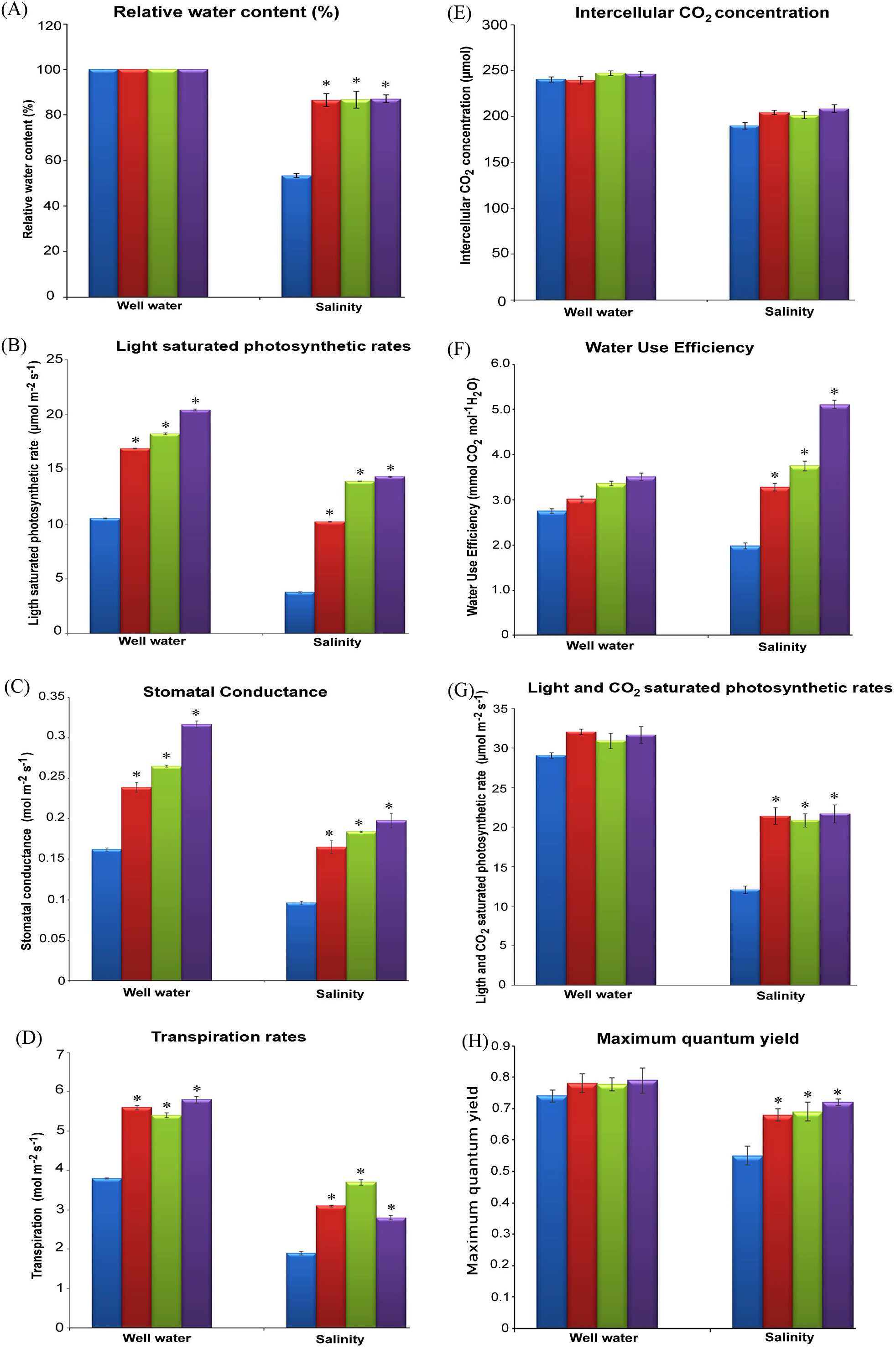
Estimation of plant water status and photosynthetic efficiencies of AT lines under salinity stress. **(A)** Leaf RWC was analysed from the fresh leaf discs sampled from AT plants subjected to well-water and 250 mM NaCl stress along with WT plants. The parameters of photosynthetic efficiency **(B)** A_Sat_; light saturated photosynthetic rates, **(C)** g_s_; somatal conductance, **(D)** E; transpiration rates, **(E)** C_i_; intercellular CO_2_ concentration, **(F)** WUE; water use efficiency, **(G)** A_MAX_; light and CO_2_ saturated photosynthetic rates and **(H)** F_V_/F_M_, maximum quantum yield were measured on well expanded, light exposed and randomly chosen leaves of AT lines and WT plants, using a portable infrared CO_2_/H_2_O gas analyzer (IRGA) LI-6400XT, under well-water and salinity stress conditions. In each treatment, 10 plants of each WT and three AT lines were used. Bar represents mean and error bar represents SE from three independent experiments. * indicates significant differences in comparison with WT at P <0.05. A10-*Ds-RFP6*, F4-*Ds-RFP31* and T8-*Ds-RFP3* are three different AT lines, WT: Wild type.

### Estimation of chlorophyll content and reducing sugars in AT lines

Under well-water conditions, no significant differences were observed for chlorophyll content and reducing sugars between WT plants and AT lines. However, under salinity stress conditions, significant increases in both chlorophyll and reducing sugars were observed in AT lines when compared to the WT plants (Fig. 4A, Fig. 4B). The chlorophyll contents of WT plants under well-water and salinity stress were 3.23 mg g^-1^ FW and 1.47 mg g^-1^ FW, respectively, while its contents in AT lines under identical conditions ranged from 3.31 to 3.58 mg g^-1^ FW and 2.91 to 3.31 mg g^-1^ FW, respectively (Fig. 4A). In addition, both WT plants and AT plants disclosed reducing sugar contents ranging from 0.521 to 0.571 mg g^-1^ FW under well-water conditions. However, imposition of salt stress caused significant increases in reducing sugar contents in WT plants (0.667 mg g^-1^ FW) as well as in AT lines (1.601 to 1.792 mg g^-1^ FW) (Fig. 4B).

**Fig. 4.**
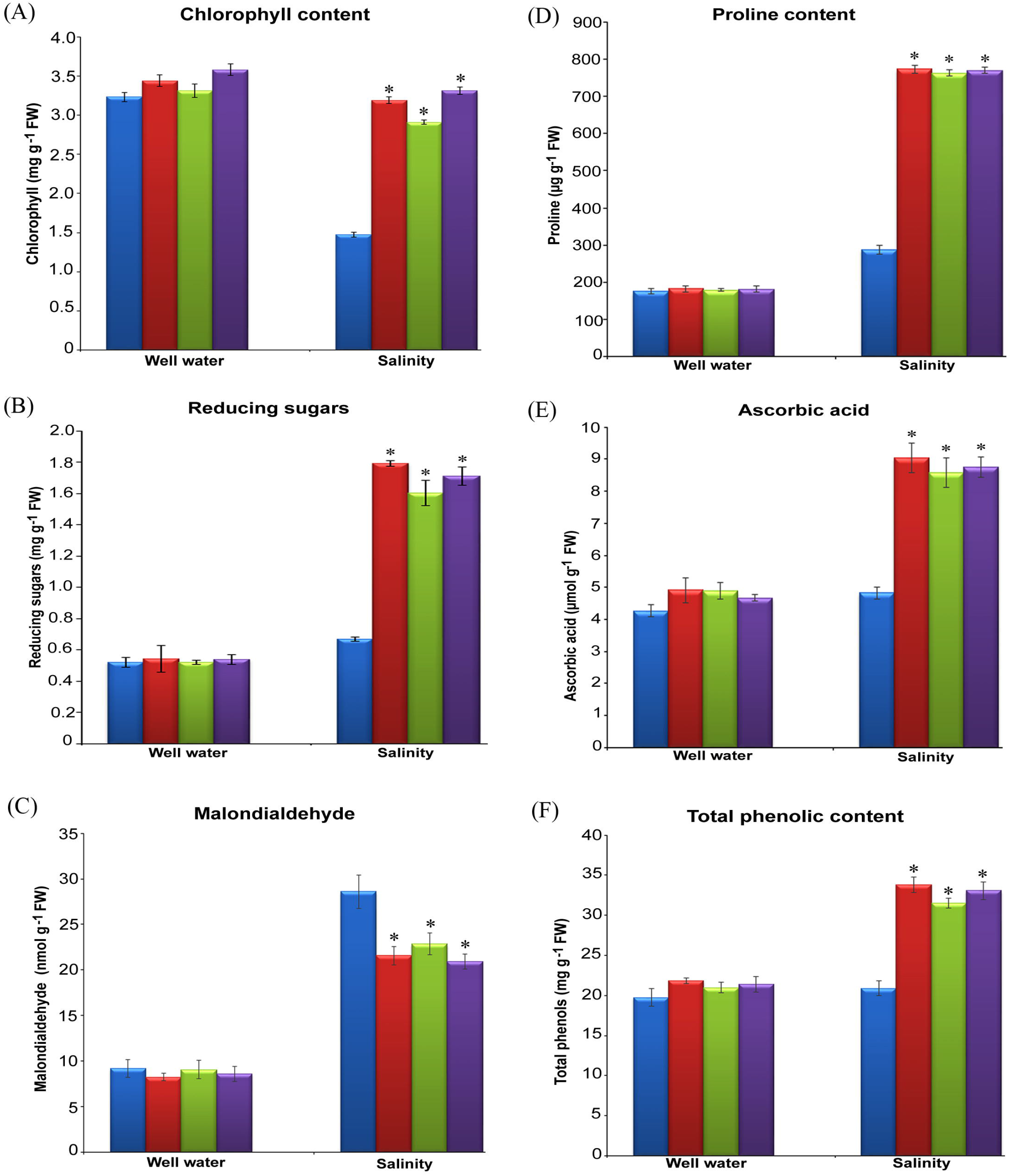
Biochemical characterization of AT lines under salinity stress conditions. **(A)** The leaf discs collected from AT lines and WT plants subjected to well-water and 250 mM NaCl stress conditions were used in estimation of total chlorophyll content. The supernatant extracted from the leaf samples of above plants were utilized in quantification of **(B)** reducing sugars, **(C)** malondialdehyde, **(D)** proline content, **(E)** ascorbic acid and **(F)** total phenolic contents. In each treatment, 10 plants of each WT and three AT lines were used. Bar represents mean and error bar represents SE from three independent experiments. * indicates significant differences in comparison with WT at P <0.05. A10-*Ds-RFP6*, F4-*Ds-RFP31* and T8-*Ds-RFP3* are three different AT lines, WT: Wild type.

### Analyses of oxidative stress parameters, osmolytes and antioxidants in AT lines

In order to assess the effects of oxidative stress, MDA content was measured in WT and AT lines under all representative conditions and no significant differences were noticed between them under well-water conditions. However, imposition of salinity stress caused significant increases in the MDA content of WT plants and AT lines, but the increases were more in WT plants when compared to AT mutants. Under salinity stress, the MDA content of WT plants was 28.58 nmol g^-1^ FW which was higher than the MDA levels recorded in the AT lines (20.67 to 22.85 μmol g^-1^ FW) (Fig. 4C). To check the osmotic adjustment ability, proline contents were estimated in WT plants and AT lines and no differences were noticed in proline levels of AT and WT plants under well-water conditions. However, the AT lines revealed significantly higher proline levels under salinity stress conditions (762 to 772 μg g^-1^ FW) when compared to the WT plants (287 μg g^-1^ FW) (Fig. 4D). Furthermore, to analyze the ROS scavenging ability, ascorbic acid (ASA) and total phenolic content (TPC) were determined in the AT lines and WT plants under salinity stress and well-water conditions. However, no significant differences were found between AT lines and WT plants in ASA and TPC contents under well-water conditions. Whereas, under salinity stress, AT lines exhibited higher ASA (8.58 to 9.04 mg g^-1^ FW) and TPC (31.52 to 33.82 mg g^-1^ FW) values when compared to WT (4.83 and 20.89 mg g^-1^ FW) plants (Fig. 4E, Fig. 4F).

### Antioxidant enzyme activities in AT lines

Under well-water conditions, no significant differences were observed in antioxidant enzyme activities of AT lines and WT plants. However, after imposition of salinity conditions, significant increases in the antioxidant enzyme activities were observed in the mutants as compared to the WT plants (Fig. 5). The catalase activity of the WT plants and AT lines varied from 0.933 to 1.086 U mg protein^-1^ min^-1^ under well-water conditions. However, salt stress could induce significant increase in the catalase activity of mutant lines (2.459 to 2.849 U mg protein^-1^ min^-1^) as compared to the low activity of WT (1.587 U mg protein^-1^ min^-1^) plants (Fig. 5A). Likewise, significant increase in the activities of SOD (15.78 to 16.46 U mg protein^-1^ min^-1^) and GPX (1.957 to 2.261 mmol min^-1^ g^-1^ FW) were recorded in AT lines, while reduced enzyme activities (8.56 U mg protein^-1^ min^-1^ and 0.814 mmol min^-1^ g^-1^ FW) were noticed in WT plants (Fig. 5B, Fig. 5C). Furthermore, marked increases in the activities of APX (3.921 to 4.182 U mg^-1^ protein) and MDHAR (44.87 to 49.25 nmol min^-1^ g^-1^ FW) and GR (1.586 to 1.829 nmol min^-1^ g^-1^ FW) were observed in AT lines compared to 2.072 U mg^-1^ protein and 18.56 nmol min^-1^ g^-1^ FW and 0.158 nmol min^-1^ g^-1^ FW in WT plants (Fig. 5D, Fig. 5E, Fig. 5F).

**Fig. 5.**
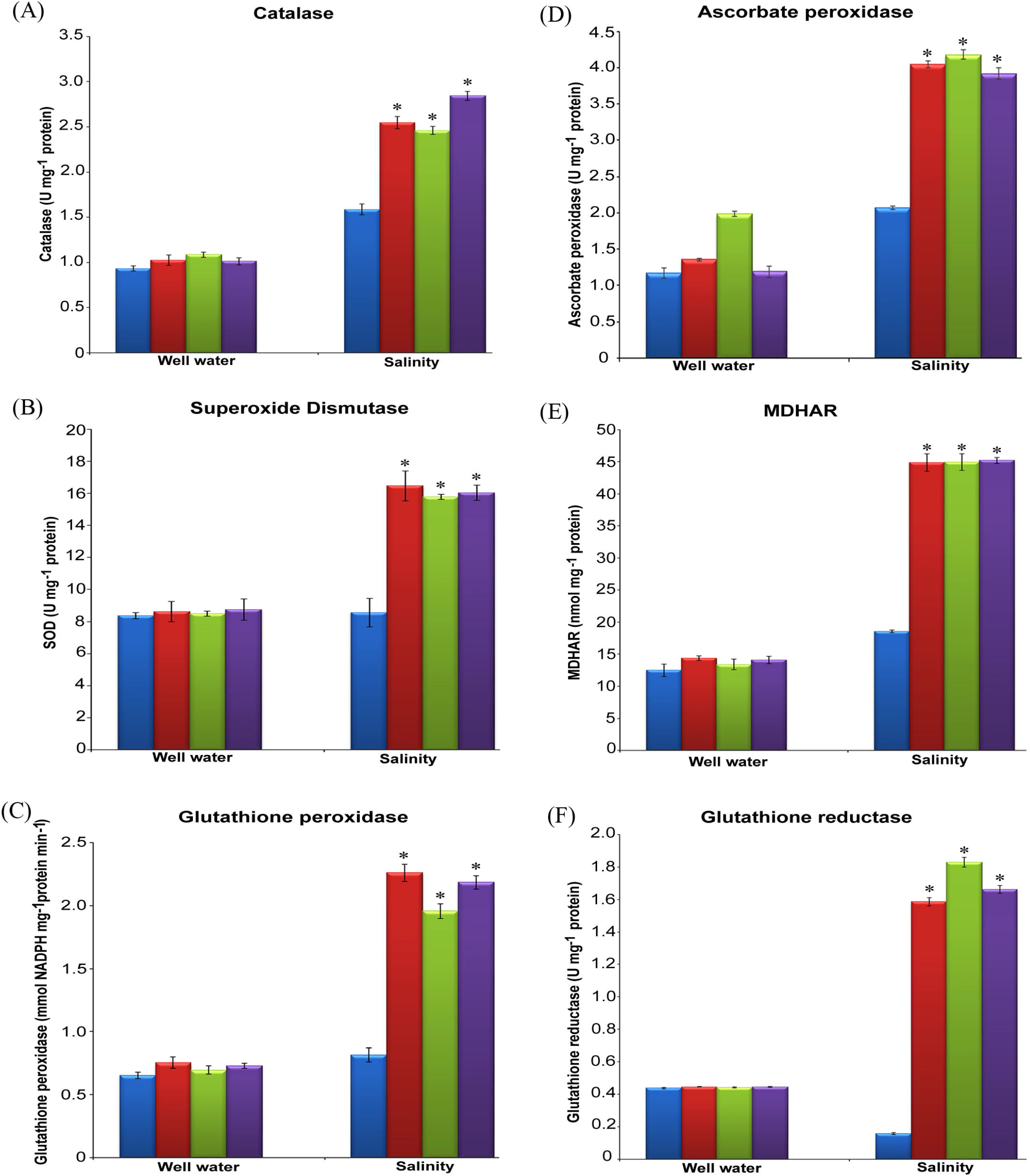
Quantification of antioxidant enzymes activities in AT lines under salinity stress conditions. AT lines and WT plants were subjected to well-water and 250 mM NaCl stress for 15 days. The supernatant extracted from the leaf material using potassium phosphate buffer were utilized for the estimation of **(A)** catalase, **(B)** superoxdie dismutase, **(C)** glutathione peroxidase, **(D)** ascorbate peroxidase, **(E)** monodehydro ascorbate reductase and (**F**) glutathione reductase. In each treatment, 10 plants of each WT and three AT lines were used. Bar represents mean and error bar represents SE from three independent experiments. * indicates significant differences in comparison with WT at P <0.05. A10-*Ds-RFP*6, F4-*Ds-RFP*31 and T8-*Ds-RFP3* are three different AT lines, WT: Wild type.

### TAIL and qRT PCR analyses of AT lines

For determining *Ds* insertion and flanking sequences in AT lines, TAIL-PCR analyses were performed using the genomic DNA isolated from A10-*Ds-RFP*6, F4-*Ds-RFP*31 and T8-*Ds-RFP*3 AT lines, using the arbitrary primer AD-2 and 3’ *Ds* specific primers (Supplementary Fig. S3). The TAIL-PCR products obtained were sequenced and analyzed using BLAST against the rice genome of Rice Annotation Project Database (RAP-DB) (http://rice.plantbiology.msu.edu/). The BLAST observations revealed that the *Ds* element is integrated at LOC_Os05g31230 (N-acetyltransferase ESCO1), LOC_Os10g36260 (expressed protein) and LOC_Os03g51080 (glutamate decarboxylase) loci in the genomes of A10-*Ds-RFP*6, F4-*Ds-RFP*31 and T8-*Ds-RFP*3 lines, respectively (Supplementary Fig. S4). Detailed sequence analyses of these mutants revealed the presence of different genes upstream and downstream (up to 10kb) at the integration site of *Ds* element in the rice genome (Table 1). qRT-PCR analyses of N-acetyltransferase ESCO1, cell division control protein 48 homolog B and acetyltransferase (GNAT family) coding genes in A10-*Ds-RFP*6 line showed 5.8-fold, 16.68-fold and 10.21-fold increases in their relative expressions as compared to WT plants (Fig. 6A). Similarly, F4-*Ds-RFP*31 line exhibited enhanced relative expression levels of 13.01-fold, 7.95-fold and 18.65-fold for mov34/MPN/PAD-1 family protein, tetratricopeptide repeat and disease resistance RPP13-like protein 1 encoding genes when compared to that of WT plant (Fig. 6B). Likewise, expression profiles of glutamate decarboxylase, inactive receptor kinase At2g26730 precursor and OsFBDUF17 - F-box and DUF domain containing protein coding genes in T8-*Ds-RFP*3 line showed 6.89-fold, 14.83-fold and 16.73-fold increases in relative expression levels as compared to WT plants grown under similar conditions (Fig. 6C). The above results clearly point out the overexpression of different genes present upstream or downstream to *Ds* element in the genomes of mutant plants.

**Fig. 6.**
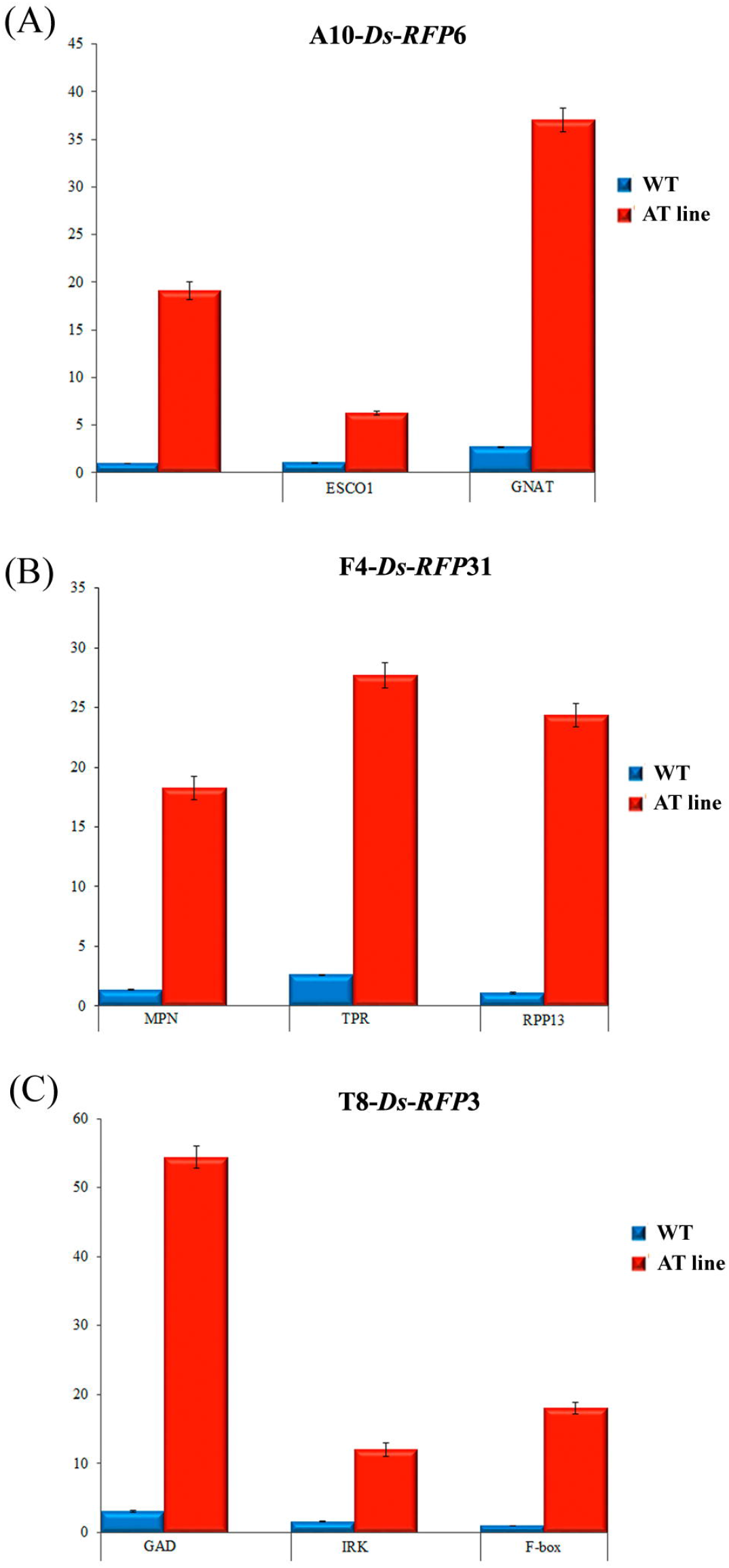
qRT-PCR analyses for the expression profiles of selected genes in AT lines of rice. Relative transcript levels of genes located upstream and downstream to *Ds* element integration site (upto 10 kb) were measured by qRT-PCR analyses. Actin was used as an internal control. The vertical column indicates the relative transcript level. Bar represents mean and error bar represents SE from three independent experiments. * indicates significant difference in comparison with WT at P <0.05. A10-*Ds-RFP*6, F4-*Ds-RFP*31 and T8-*Ds-RFP*3 are three different AT lines, WT: Wild type.

**Table 1.**
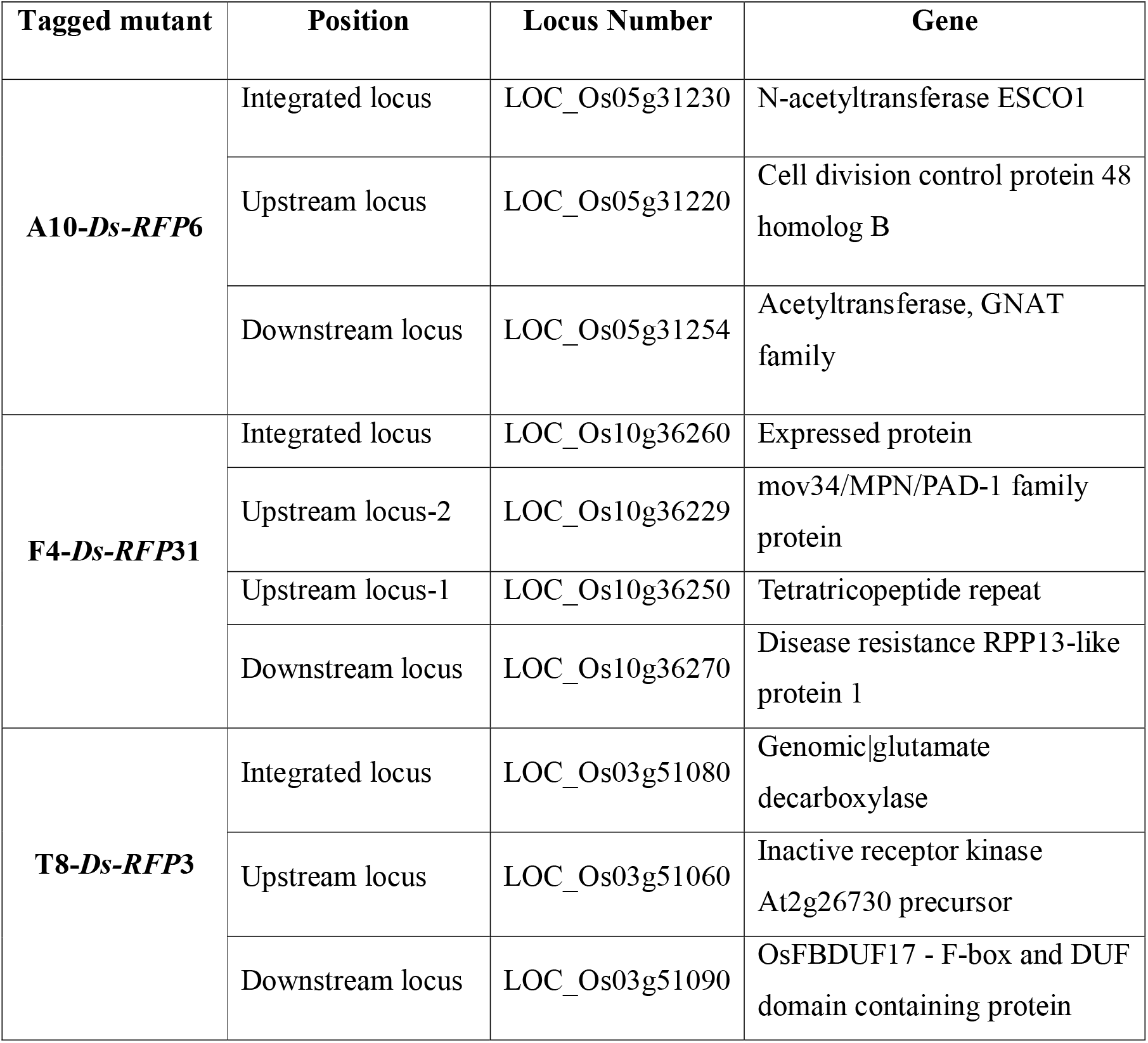
Location of *Ds* element and upstream & downstream genes at the *Ds* integrated site in the genomes of AT rice lines

| Tagged mutant | Position | Locus Number | Gene |
| --- | --- | --- | --- |
| **A10-*Ds*-RFP6** | Integrated locus | LOC_Os05g31230 | N-acetyltransferase ESCO1 |
|  | Upstream locus | LOC_Os05g31220 | Cell division control protein 48 homolog B |
|  | Downstream locus | LOC_Os05g31254 | Acetyltransferase, GNAT family |
| **F4-*Ds*-RFP31** | Integrated locus | LOC_Os10g36260 | Expressed protein |
|  | Upstream locus-2 | LOC_Os10g36229 | mov34/MPN/PAD-1 family protein |
|  | Upstream locus-1 | LOC_Os10g36250 | Tetratricopeptide repeat |
|  | Downstream locus | LOC_Os10g36270 | Disease resistance RPP13-like protein 1 |
| **T8-*Ds*-RFP3** | Integrated locus | LOC_Os03g51080 | Genomic\|glutamate decarboxylase |
|  | Upstream locus | LOC_Os03g51060 | Inactive receptor kinase At2g26730 precursor |
|  | Downstream locus | LOC_Os03g51090 | OsFBDUF17 - F-box and DUF domain containing protein |

## Discussion

Most often, loss-of-function mutagenesis may not lead to identifiable phenotypes owing to functional redundancy of genes and sometimes loss-of-function mutations might prove lethal for certain genes. However, adoption of gain-of-function mutagenesis might overcome these limitations, and in this context, activation tagging proved to be an effective method to generate gain-of-function mutants with improved agronomic traits (Weigel *et al*., 2000). In the present study, analyses of three gain of function AT lines revealed significant increases in growth, development and yield potential, both under well-water and salinity stress conditions when compared to their WT counterparts. The AT rice lines exhibited higher seed germination rates under salinity stress conditions which got translated into better plant growth and development at vegetative and reproductive stages, in comparison to the WT plants, owing to their enhanced tolerance to salinity stress. Increased seed germination rates and improved seedling establishment potential under adverse conditions revealed significant association with improved productivity in several crop plants (Hamayun *et al*., 2015; Shu *et al*., 2017; Shahzad *et al*., 2019). Furthermore, a stringent positive correlation was observed between seed germination rate and post germination effects under prevailing unfavourable conditions owing to rapid acclimatization to abnormal stress conditions (Donohue *et al*., 2010; Shahzad *et al*., 2019).

In the present investigation, AT lines disclosed higher RWC levels under salinity stress when compared to the WT plants, plausibly, owing to their higher plant water retention status and delayed onset of stress symptoms. The onset of stress related symptoms in plants grown under different abiotic stress conditions were found to be associated with the plant water status. Alterations in the leaf RWC could serve as a reliable parameter to assess the plant water status (Reddy *et al*., 2017; Sekhar *et al*., 2017). Under salinity stress, AT rice lines exhibited increased photosynthetic performance translating into higher yields owing to their superior plant water status than that of WT plants. Earlier, strong positive correlations were observed between RWC levels and photosynthetic efficiency, resulting in increased yield potential under salinity stress conditions (Nxele *et al*., 2017; Rahneshan *et al*., 2018). Previous observations have indicated that fluctuations in crop yields were proportionally related to variations in photosynthetic efficiency under normal and stressed conditions which were estimated in relation to alterations in light saturated photosynthetic rates (A_sat_) (Sekhar *et al*., 2020). The AT lines exhibited higher A_sat_ under well-water conditions indicating improved ribulose 1,5-bis phosphate (RuBP) carboxylation and translating into increased yields, when compared to WT plants. Moreover, imposition of salinity stress could affect photosynthetic performance of crops by decreasing A_sat_ values and favoring RuBP oxygenation over carboxylation, resulting in declined plant growth and yields (Chaves *et al*., 2009).

Reduced photosynthetic efficiency of crops, under various stress conditions including salinity, was primarily linked to stomatal conductance (g_s_) limitations by stomatal closure via-ABA signalling mechanism, causing reduced A_Sat_ (Flexas *et al*., 2008; Lawlor and Tezra, 2009; Gupta and Huang, 2014). Whereas, AT rice lines revealed significant higher A_sat_ under salinity stress, when compared to WT plants, plausibly owing to increased Rubisco carboxylation. Moreover, the enhanced photosynthetic performance of AT lines under salinity stress, is mainly attributable to greater g_s_ and higher C;, favouring CO_2_/O_2_ ratio in the vicinity of Rubisco active site, and minimal stomatal limitations when compared to WT plants. By contrast, WT plants disclosed early stomatal closure under salinity stress and restricted entry of CO_2_ leading to decreased photosynthetic efficiency, eventually resulting in declined growth and lower yields. Moreover, AT lines revealed higher E values both under well-water and salinity stress conditions, plausibly, owing to linkage between increased g_s_ and greater transpiration pull, leading to higher plant water status and delayed stress related symptoms, as compared to WT plants. Most often, stringent positive correlations were observed in plants between changes in g_s_ and E values when grown under well-water and stress conditions (Reddy *et al*., 2019; Sreeharsha *et al*., 2019).

AT lines exhibited increased WUE_i_, compared to the WT plants, owing to higher A_sat_ and its association with increments in biomass yields both under well-water and salinity stress conditions mainly due to their superior photosynthetic efficiency. Furthermore, improved WUE_i_, caused by reduced E and/or greater A_sat_, delayed stress related symptoms, and enhanced photosynthetic efficiency, resulted in improved crop yields (Reddy *et al*., 2017; Sekhar *et al*., 2017). In the present study, AT plants revealed higher AMax both under wellwater and salinity conditions, suggesting that their improved adaptability contributed to increased productivity under elevated atmospheric CO_2_ even when grown under salinity stress. Earlier studies indicated that crops with augmented light and CO_2_ saturated photosynthetic rates (A_Max_) demonstrated superior photosynthetic efficiency under predicted future atmospheric CO_2_ conditions (Rosenthal *et al*., 2011).

Moreover, alterations in photosystem II (PS-II) efficiency could proportionately influence A_sat_, since the rate-limiting enzymes of Calvin-Benson cycle require both ATP and reduced nicotinamide adenine dinucleotide phosphate (NADPH) for their optimum activity (Sekhar *et al*., 2014, 2015). Additionally, salinity stress was found to reduce PS-II efficiency by hampering electron flow from PS-II to PS-I, resulting in diminished production of ATP and NADPH, thereby reducing photosynthetic performance and productivity (Kalaji *et al*., 2011; Shu *et al*., 2012; Pan *et al*., 2020). However, the AT lines exhibited higher F_V_/F_M_, suggesting improved PS-II efficiency both under well-water and salinity conditions, thus contributing substantially to photosynthetic efficiency and yield components when compared with WT plants. The above observations amply indicate that AT plants could minimize both stomatal and non-stomatal limitations under salinity stress conditions, and thereby enhance photosynthesis resulting in increased yield components. In comparison, WT plants failed to perform on par with the AT mutants under stress conditions.

In the present study, imposition of salinity stress caused significant increases in the levels of ROS scavenging enzymes of both AT lines and WT plants owing to the activation of ROS signalling mechanism. However, AT plants disclosed greater activities of enzymes belonging to antioxidant metabolism such as peroxidase cycle (SOD, Catalase and GPX) and enzymes of ascorbate-glutathione cycle (ASC-GSH), namely APX, GR and MDHAR causing reduced oxidative damage and delayed stress symptoms when compared to WT plants. Apart from improved photosynthesis, plants could delay various abiotic stress symptoms either by minimizing the ROS production and/or reduced ROS accumulation owing to presence of enzymatic as well as non-enzymatic antioxidant systems (Gill and Tuteja, 2010; AbdElgawad *et al*., 2016; Al Hassan *et al*., 2017; Kibria *et al*., 2017).

AT lines exhibited increased accumulation of proline under salinity which was found associated with higher levels of stress tolerance compared to their WT counterparts. Salinity was reported to induce accumulation of proline, a major stress induced osmolyte, and its concentration was found to be directly proportional with the plant stress tolerance capacity (Hayat *et al*., 2012; Huang *et al*., 2013). AT lines exhibited elevated levels of reducing sugars under salinity stress which were found associated with increased water retention capacity and elevated plant water status as revealed by higher RWC, resulting in mitigated stress symptoms. Plants grown under different abiotic stress conditions developed adaptability to conserve more water by accumulating soluble sugars as compatible osmolytes, leading to higher plant water status and reduced stress symptoms (Yin *et al*., 2010; Sami *et al*., 2016; Rahneshan *et al*., 2018). AT lines disclosed increased concentration of ASA and polyphenols during salinity stress, as compared to WT plants, owing to their effective nonenzymatic antioxidant system promoting ROS detoxification. Imposition of different abiotic stresses triggered increases in the accumulation of antioxidant compounds of ASA and polyphenols, resulting in incessant ROS detoxification (Ghasemzadeh *et al*., 2010; Al Kharusi *et al*., 2019). In addition, stringent positive correlations were observed between changes in antioxidant enzyme activities (ASA-GSH cycle) and ASA at various environmental conditions (Gill & Tuteja, 2010; AbdElgawad *et al*., 2016). In this study, reduced MDA content observed in AT lines as compared to WT plants, obviated the e ective detoxification of ROS and declined oxidative lipid injury in plant cells. Moreover, delayed senescence and/or severe stress symptoms in AT lines during salinity stress could be associated with increased ROS scavenging culminating in reduced oxidative damage and enhanced yields. Furthermore, salinity stress caused loss of cellular homeostasis and excessive generation of ROS leading to lipid membrane peroxidation, resulting in significant increases in MDA content (Akcay *et al*., 2012; Rangani *et al*., 2016).

AT plants of A10-*DS-RFP6* exhibited substantial 16.68-fold increase in CDC48 expression levels in comparison to WT plants. It was reported that the CDC48 could play a crucial regulatory role in plant development besides protein degradation through ubiquitin-proteasome system (UPS) and endoplasmic reticulum-associated protein degradation (ERAD) system (Begue *et al*., 2019). Shi *et al*., (2019) observed that the transgenic rice overexpressing *Os*CDC48 promoted increased tiller number and grain yield. Based on present results, we postulate that CDC48 plausibly contributes to the maintenance of cellular homeostasis by mediating multiple regulatory processes. Moreover, elevated expression levels of N-acetyltransferase ESCO1 and GNAT-acetyltransferases in the AT plants implicate their intrinsic role in the regulated expression of stress responsive genes. Kalamaki *et al*., (2009) reported the involvement of acetyltransferases of GNAT family in conferring tolerance to drought, salinity and cold by controlling the expressions of diverse stress responsive genes.

Elevated relative expression levels of mov34/MPN/PAD-1 family gene, tetratricopeptide repeat containing gene and disease resistant RPP13-like protein 1 in F4-*Ds-RFP*31 AT line suggest the involvement of these genes in various biological processes including abiotic tolerance in rice. Schaper & Anisimova (2016) reported that plant proteins containing conserved tetratricopeptide repeats (TPRs) were involved in both primary metabolism (cellular biosynthetic processes) and secondary metabolism (response to abiotic stimuli). Higher expression levels of inactive receptor kinase At2g26730 precursor, glutamate decarboxylase and *Os*FBDUF17-F-box and DUF domain containing genes of T8-*Ds-RFP3* AT plants indicate their probable involvement in various aspects of plant growth, development and response to different external stimuli. Especially, increased expression of *Os*FBDUF17 - F-box and DUF domain containing genes in the AT line could be attributed to its enhanced tolerance levels to salinity stress conditions. In wheat, it was found that increased glutamate decarboxylase expression and accumulation of GABA were associated with higher levels of tolerance to salinity and osmotic stress (AL-Quraan *et al*., 2013).

To sum up, an overview of present results dealing with selected rice AT lines generated by employing the versatile maize Ac/*Ds* system, demonstrated rare improvements in various multigenic morphometric characters, besides exhibiting enhanced tolerance to salinity stress at different stages of growth and development. Furthermore, qRT-PCR analyses of AT lines revealed remarkable expression levels of several candidate genes responsible for synthesis of CDC48, N-acetyltransferases, receptor kinase, glutamate decarboxylase and F-box-DUF proteins involved in various aspects of plant growth and differentiation. Hopefully, analyses of these genes based on their overexpression and gene silencing approaches could provide further insights into their functionality and involvement in contributing to improvements of different agronomic traits of rice. Furthermore, the promising AT rice lines plausibly serve as a key genetic resource to accomplish strategic improvements in major yield components of elite rice cultivars besides evolvement of novel genotypes bestowed with genetic resilience to cope with the challenges of rapid climatechange conditions.

## Acknowledgments

This work was supported by a grant (BT/PR-13105/AGR/02/684/2009) from the Department of Biotechnology, Government of India. Authors are grateful to Prof. Venkatesan Sundaresan, University of California, Davis, USA, for providing pSQ5 activation tagging vector. KVR is thankful to the Council of Scientific and Industrial Research, New Delhi and KMS is thankful to University Grants Commission, DSKPDF, New Delhi, for the award of Research Fellowships. We thank Prof. S. Rajagopal, Department of Plant Sciences, University of Hyderabad, for allowing us to use glass house facilities. We are grateful to Prof. T. Papi Reddy for the critical reading and improving the manuscript.

## Author Contributions

Conceived and designed the experiments: KVR, VDR, ARR. Performed the experiments: KVR, KMS. Analyzed the data: KVR, VDR, ARR, KVR, KMS. Wrote the paper: KVR, KVR, KMS.

## Supporting Information

**Supplementary Fig. S1** PCR analysis of AT lines for the presence of *RFP* region using *RFP* forward and *nos* reverse primers

**Supplementary Fig. S2** PCR analysis of AT lines for the presence of 4X enhancer region using *En* forward and *En* reverse primers

**Supplementary Fig. S3** TAIL-PCR analyses of AT lines of rice

**Supplementary Fig. S4** BLAST analysis of AT rice lines showing integration site of *Ds* element and genes located on either side of *Ds* in the rice genome

**Supplementary Table S1** Thermal cycling conditions employed for primary, secondary and tertiary TAIL-PCR using AD and 3 □ *Ds* primers

**Supplementary Table S2** Primers employed for qRT-PCR analyses of AT rice lines

